# Developmental trajectories of cognitive traits in meerkats match socio-ecological demands

**DOI:** 10.1101/2025.10.29.685340

**Authors:** Tommaso Saccà, Elisa Protopapa, Adrian V. Jaeggi, Sofia I. F. Forss

## Abstract

Understanding cognitive development is fundamental for explaining variation in adult cognitive phenotypes, and thus the processes driving cognitive evolution. The ontogeny of cognitive traits is likely influenced by both population-wide pressures, such as ecological demands, and individual-specific factors, including early life experiences. To properly investigate cognitive variation, we must therefore identify species-level developmental trajectories and individual variation from normative ontogeny. We studied the ontogeny of three cognitive traits—inhibition control, spatial cognition, and physical problem-solving skills—in 28 wild meerkats (*Suricata suricatta*) from eight litters (seven groups). Our longitudinal study followed individuals from early life to nutritional independence and into sub-adulthood. We found that rates of development (learning curves) varied among traits, reflecting the timing of socio-ecological pressures in this species. Performance of cognitive traits did not correlate over ontogeny nor at specific time points, suggesting independent processes underlying each measured trait. While the development of inhibitory control appeared conserved, that of spatial cognition and physical problem-solving showed substantive individual differences. Furthermore, only physical problem-solving showed consistency in performances over time, emerging near nutritional independence, reflecting its ecological importance early in life. Future research should determine the drivers of individual developmental variation and how such differences translate into fitness consequences.

## Introduction

Individual differences are increasingly relevant for explaining cognitive evolution (1). An important and underappreciated source of individual differences in cognition is developmental effects (2). Despite some recent work (3–5), this research is still in its infancy and studies investigating the ontogeny of cognitive traits remain especially sparse beyond primates and in wild conditions. Individual variation in cognitive developmental trajectories could facilitate adaptive responses when new selective pressures on existing phenotypes arise due to environmental variability (6–8). For example, given that advanced cognitive abilities may require larger brains, which are energetically costly (9) and thus trade-off with other functions such as reproduction (10,11), periods of food abundance may promote faster cognitive development while periods of food scarcity could select for slower developmental rates (12,13). When experienced during early life, ecological pressures such as diet (14), physical or social environment (3,15–17), have long lasting impacts on the adult cognitive phenotype. To explain how early-life pressures shape individual-level cognitive development, we need a clear account of population-level developmental trajectories. Crucially, the study of phylogenetic differences in developmental trajectories, known as heterochrony, has delivered important scientific advancements in our understanding of cognitive evolution (18–21). For example, comparative developmental studies of face recognition, gaze following, and goal-directed actions suggest that humans rely more heavily on social information than any of our closest primate relatives, with such differences emerging early in ontogeny (18,22). Furthermore, chimpanzees (*Pan troglodytes*) develop both spatial cognition and inhibitory control more rapidly than bonobos (*Pan paniscus*), which arguably reflects differences in their feeding ecology and social system (23,24).

Here we present a detailed, longitudinal investigation of early cognitive development in a wild social carnivorous mammal, the meerkat (*Suricata suricatta*). We studied meerkats throughout their ontogeny, from nutritional dependence to sub-adulthood, by experimentally measuring the developmental trajectories of three distinct physical cognitive skills: inhibitory control, spatial cognition, and extractive foraging represented through a physical problem-solving task (Figure 1). These cognitive traits reflect pressures experienced by meerkats in their ecological and social environment. Meerkats are highly social, cooperatively-breeding mammals in which adults and sub-adults contribute to raising offspring, defending against predators and outgroup conspecifics, and burrow maintenance (25). Cooperative breeding has been proposed to select for enhanced executive functions, including inhibitory control, because individuals must supress self-serving behaviours to engage in cooperative activities (26,27), but see (28). Moreover, meerkats live in environments characterised by high predation pressure and limited physical refuges (i.e., boltholes) from aerial predators (29). This makes the development of spatial cognition particularly important as immature individuals must learn proper, adult-like responses to alarm calls (30) and acquire adult-level knowledge of bolthole locations (31). Finally, the development of physical problem-solving is ecologically relevant because meerkats inhabit a complex foraging niche characterised by seasonality and habitat heterogeneity (32,33), and immature individuals rely on adults to learn foraging skills, define their food repertoire and acquire complex prey-handling skills (34–36). We would also expect meerkat life-history to influence the cognitive performance of males and females. In meerkats, where reproduction is largely monopolised by dominant individuals (29), females are philopatric and can maximise fitness by attaining dominance within their birth group (37). Males in contrast must disperse to secure a dominance position, and as subordinates most of their reproductive success comes from prospecting and gaining extra-group copulations (29,38,39). Dispersal is risky due to predators and aggression from neighbouring groups (29), and its success likely involves the ability to navigate unfamiliar environments (spatial cognition), to efficiently locate food (physical problem-solving), and to prioritize effective decisions over prepotent responses (inhibition control) in this dangerous, highly arousing context. Overall, success in dispersal events requires effective decision making under limited cues and has been proposed to contribute to socially driven cognitive evolution (40).

**Figure 1:**
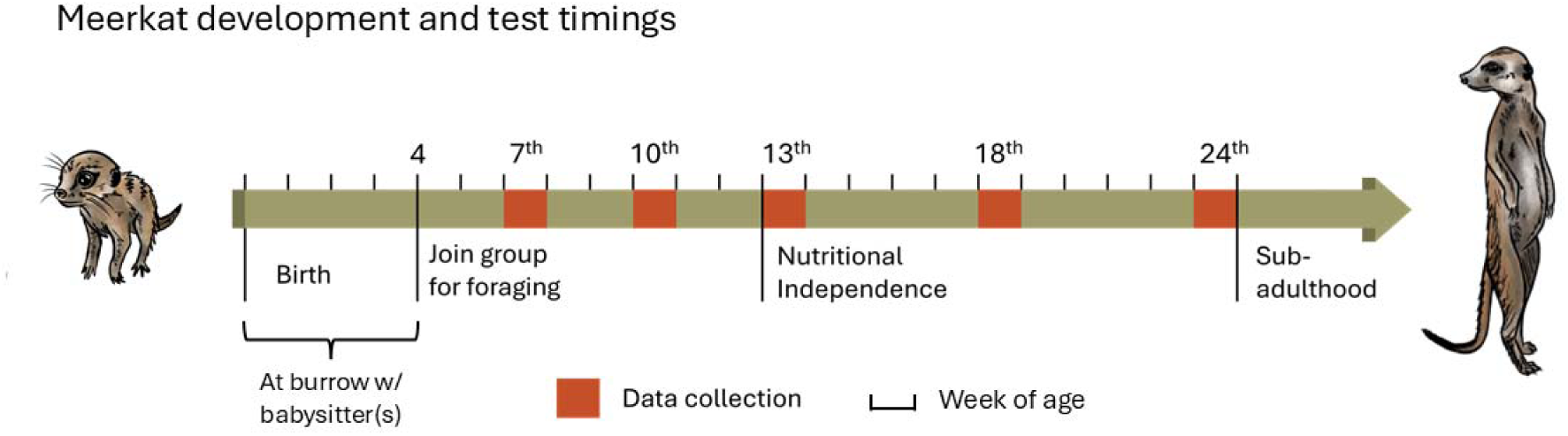
Overview of meerkat early life development and data collection time points. Weeks of age are indicated as black ticks, testing weeks as orange rectangles, and key life history stages with longer ticks (see Methods for details).

Altogether, meerkat pups depend on adults to develop essential survival skills, such as foraging and anti-predator behaviours (41). Ontogenetic reliance on social inputs has been linked to predictions about domain-general cognition (i.e. correlated performance across cognitive traits). Extensive reliance on social learning has been argued to favour the emergence of human-like domain-general intelligence, by enhancing learning efficiency and capacity during development which translate across cognitive domains (the Cultural Intelligence Hypothesis) (42). On the other hand, strong reliance on social learning may reduce selection pressure on independent, generalized learning abilities, and avoid the associated energetic costs of large brains (43). In addition, it has been proposed that cooperative breeding systems present unique challenges that select for enhanced socio-cognitive abilities (the Cooperative Breeding Hypothesis) (26,44,45), as well as reducing energetic constrains on larger brains (46). Therefore, the social ecology and cooperative breeding system of meerkats provide a unique opportunity to evaluate hypotheses on the evolution of cognition in a non-primate mammalian species.

In this study we repeatedly assessed individual meerkat pups over multiple key developmental life stages (Figure 1). Our longitudinal approach enabled us to ask I) whether distinct cognitive traits develop in parallel within individuals, II) whether relationships among traits exist and whether these change over ontogeny, and lastly III) whether performance was consistent across developmental stages. Finally, we discuss population-level developmental trajectories for each trait in the context of species-specific socio-ecological demands.

## Methods

### Study population

We conducted our study at the Kalahari Research Centre, located in the Kuruman River Reserve (KRR) (26°59′ S, 21°50′ E) in the Kalahari Desert in South Africa. The KRR is a semi-arid savanna with distinct macro-habitats such as red sand dune and shrub grassland (33). Vegetation is sparse during the dry season and more abundant during the wet season (47). Rainfall averages 266mm per year, and temperatures are on average 30.3 ^○^C during the wet season (maximum 44.2 ^○^C) and 10.1 ^○^C during the dry season (minimum -11.9 ^○^C) (33). Here, we investigated 28 wild meerkats (12 females and 16 males), from seven groups and eight litters with average 3.5 pups (range 2 to 4) over two wet Seasons (December 2022 – May 2023, and November 2023 – May 2024). All groups and individuals included in our study were habituated to close observation.

### The longitudinal developmental approach

For the first three weeks of life, meerkat pups stay below ground, after which they emerge and remain close to the natal burrow. During this time, they are always joined by usually at least one older group member to ensure their safety (48). At about four weeks of age, they start to leave the natal burrow and join their group foraging. They rely on adults for food acquisition until about 12 weeks of age, after which pups are considered nutritionally independent (49). They are considered sub-adult at 24 weeks of age, when they first start to participate in all cooperative activities, and adults at one year of age after reaching physiological reproductive maturity (50). We tested individuals on their cognitive performance at five key points during their development (see Figure 1). The earliest point was the 7^th^ week of life, as this was when meerkat pups would first show interest in interacting with cognitive tests. At this time pups had left the safety of their natal burrow and were consistently joining their group foraging already for about 2-3 weeks. We re-tested all pups at 10, 13, 18 and 24 weeks to evenly space sampling effort around nutritional independence, with two measurements before, one during, and two after, with the final point corresponding to early sub-adulthood. Testing weeks were based on supposed date of birth and we allowed on average a two-day flexibility for testing individuals before and after the end of the formal week for the 7^th^, 10^th^ and 13^th^ week, and four-day flexibility for the 18^th^ and 24^th^ week, to reduce logistical problems in data collection. At times, we could not test individuals due to external limitations or to individuals not being interested in the test apparatus. In other instances, we excluded single tests post hoc for not meeting sufficient test criteria. A complete list of deviations from planned data collection can be found in Supplementary Table S1. Differences in the number of presentations across developmental weeks were accounted for in the analysis.

## Test paradigms of cognitive traits

### Inhibition control task

To test for the ability to inhibit incorrect intuitive responses, we used a standardized inhibitory control task (sometimes referred to as the “cylinder test” (51)), which in our study consisted of a rectangular box with open ends containing a food reward in the middle. Prior to testing, at each developmental week, subjects were habituated to an opaque version of the test apparatus that was presented with the opaque barrier facing the subject. Before testing, we ensured that each subject had learnt to disregard the barrier and access the reward from the open sides, after which they were considered ready to be tested on their ability to recall the learned response and inhibit prepotent but incorrect responses. Detailed information about the habituation procedure can be found in Supplementary Text S2. During testing, we presented five times per developmental week to each subject a transparent version of the apparatus with the transparent barrier facing the subject (Figure 2). We considered a trial successful if the subject did not touch the transparent side before going around the open side and accessing the food reward.

**Figure 2:**
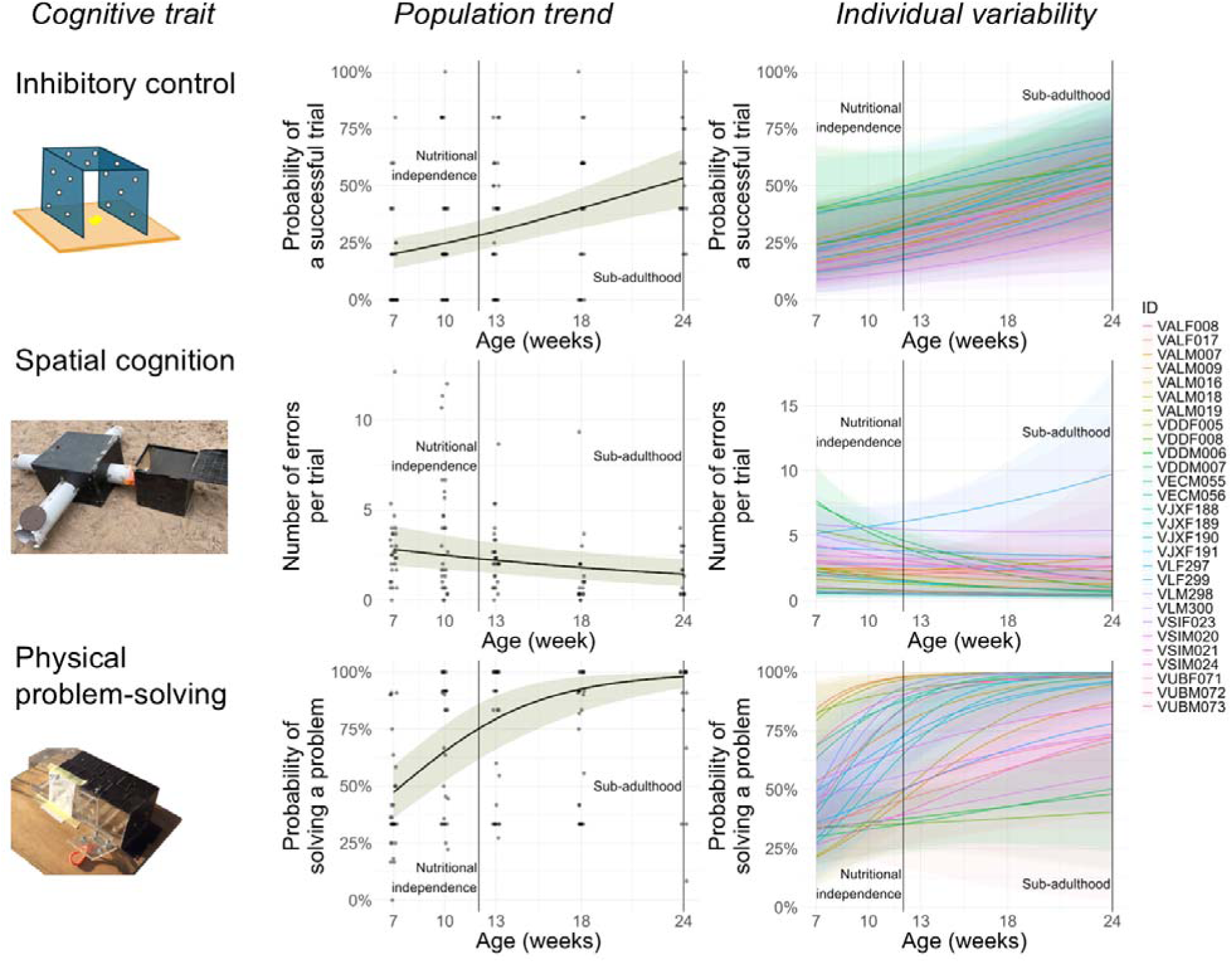
Population-level developmental trajectories (second column) and individual ontogenetic variability (third column) for three cognitive tasks (first column). Vertical lines indicate nutritional independence and sub-adulthood. In all plots, age (developmental week) is on the x-axis and task performance on the y-axis. Population-level trajectories: black line shows the modelled population mean, green shading indicates the 95% confidence interval, and observed data are plotted in the background. Data points are semi-transparent, with darker shades indicating overlapping points, and jittered along the x-axis for clarity. Individual variability: lines show modelled trajectories for each individual, with shaded 95% confidence intervals; darker shading indicates overlapping intervals. For population-level spatial cognition, predictions average across seasons, and all population-level trends are averaged across sexes.

### Spatial cognition task

We evaluated spatial cognition using a four-armed maze with a central chamber and a starting box at the end of one arm. The starting box, the central chamber and all arms were fully opaque (Figure 2), while the top of the central chamber and the ends of each arm varied between data collection seasons (transparent in the first season and opaque in the second). This change was done to further minimize subjects’ possible reliance on external cues that could not be kept constant between trials. Subjects were habituated to the apparatus each developmental week before testing. Additional details about habituation can be found in Supplementary Text S2. During testing, three exits were covered by plexiglass, while the fourth was always open and connected to the starting box. One of the covered exits could be opened, constituting the correct exit, while the others two were fastened shut. During testing, we assessed stress levels by monitoring characteristic lost calls (52) and other vocalizations. If subjects showed even mild signs of distress, judged from a combination of vocalizations and physical cues of arousal, we guided them to the correct exit by inserting our hand there and providing visual and auditory cues. We considered the first successful trial as the first trial where the individual successfully found the correct exit without external help. From then on, we tested each subject in three more trials. In those instances, if the subject became too stressed, we removed it from the test without guiding it to the correct exit and we considered the trial a mistrial.

The design of the spatial cognition task slightly differed between the first and second season of data collection. During the first season the plastic ends of the three exits and the top of the central chamber were made of transparent plexiglass. In addition, the correct exit for each subject could either be the same as the previous week or change. During the seventh week of the first study group, the main chamber of the maze was fully transparent. This data point was considered as the same to the datapoints from the first season, for sample size limitations on this design. During the second season the plastic end of the three exits were opaque, and the correct exit for each subject always changed between successive weeks, to avoid a purely associative learning effect.

### Physical problem-solving task

To test the ability to solve physical problems to access a food reward, we presented an apparatus with four chambers, each containing a food reward inside (see Figure 2). Three chambers were covered by a different obstacle (henceforth: ‘problem’) to be removed or opened, with one chamber left unobstructed to maintain motivation to participate. Each problem could be solved with physical motions routinely used while foraging, specifically digging, poking, and pulling. Before each week of testing subjects were habituated to the test apparatus to avoid any potential effects linked to neophobia and approaching of novel human-made artefacts (53). Further details about habituation procedure and apparatus design can be found in Supplementary Text S2. During testing, we presented the test apparatus to each subject four times during each developmental week. We considered a problem solved if the subject had found a way to access and eat the food reward inside, even if we could not infer whether the action used was intended to solve the problem. We did not consider a chamber solved if the problem was opened by accident by an involuntary bump or any other sort of non-active solution.

#### Role of memory

To investigate whether the ontogeny of performance in the problem-solving task was primarily influenced by learning and memory from repeated exposure over development, we presented the apparatus to an additional six subjects (four males, two females) at ten weeks of age who had not encountered it before at seven weeks of age and recorded their performance. We hypothesised that, if the increase in performance was mainly due to memory and prior exposure, these six subjects would (a) perform similarly to subjects first presented with the apparatus at seven weeks of age, and (b) perform worse than those same subjects when the latter were re-presented with the apparatus at ten weeks of age.

## Data scoring and statistical analyses

We recorded each presentation using video cameras (Hero 9 Black: GoPro Inc., 2020), and we scored cognitive task performances using the software Boris (54) and a predefined ethogram (Supplementary Table S3). For the spatial cognition test, to account for the effect of potential stress on performance, we noted as a binary variable for each trial if the subject produced any “lost call” vocalization during testing. For the physical problem-solving test, to account for motivational levels, which is known to influence cognitive performance (55), we scored the time the subject was interacting with or investigating the test apparatus. In addition, to account for differences in study design, we noted down whether the design was linked to the first or second season of testing. All analyses were conducted R v4.4.0 (56), using RStudio as an interface (57). For Bayesian hierarchical and multivariate model fitting we used the package BRMS (58). Throughout our analyses, we used weakly informative priors (intercepts: normal(0,2); fixed effects: normal(0,2); standard deviations of random effects: exponential(2); correlations of random effects: LKJ(2)) to aid model convergence and regularize parameter estimates, thus reducing type I errors (59). For all models, we confirmed proper model fitting by visual inspection of chain mixing, Rhat values for convergence (all = 1.00) and effective sample size measures (all > 1000).

### Population-level developmental trajectories and individual variation

To investigate individual performance over development for each cognitive trait we fitted three sets of Generalized Linear Mixed Models (GLMM) in a Bayesian framework, one for each cognitive task. We added to each of the three models a random individual intercept and a random individual slope for the effect of developmental week, treated as a continuous variable (henceforth: week). This allowed the models to represent the developmental trend of each subject, accounting for both individual baseline performance after adjusting for the effect of covariates (individual random intercept), and individual differences in developmental rates (individual random slope for the effect of week on task performance). These random effects were estimated as individual deviations (positive or negative) relative to, respectively, the population-level performance (global intercept) and developmental trend (fixed effect of week). This structure also enabled us to investigate to which extent performances could be attributed to individual characteristics. To do so, we calculated the proportion of variance explained by the random intercepts and slopes not explained by fixed effects using the function “variance_decomposition” from the package Performance (60). This is the suggested approach when working with non-Guassian distribution and complex random structures in Bayesian models fitted with BRMS package. This approach is functionally equivalent to the widely used “adjusted intraclass correlation coefficient” (adjusted ICC).

### Inhibition control

For the performance in the inhibitory control task we used as the response variable the number of trials passed versus the number of total trials per week of testing, assuming a binomial distribution, and we added as explanatory variable the effect of week and of sex.

### Spatial cognition

For the performance in the spatial cognition task, we fitted two models, to investigate competing variables linked to study design. First, we used the total amount of errors over the number of trials per week of testing as a response variable, assuming a Poisson distribution, including as covariates (a) the number of trials per week where the subject showed mild signs of distress, (b) week, (c) sex, and (d) a binary variable for whether the correct exit was kept the same as for the previous week of testing. Secondly, we fitted a model where all variables were equivalent, but we excluded the binary variable for whether the correct exit changed from the previous week. Instead, we fitted a variable for study design, distinguishing between first and second season. In both models, for the total amount of errors and number of trials we excluded the performance in the first trial per week, since we could not assume the subjects knew the solution and the performance was more due to chance than actual cognitive processing.

### Stress and season of testing

We investigated whether the physical design of the spatial cognition test (i.e., ceiling and arm openings transparent during the first season and opaque during the second season) influenced the level of stress experienced by subjects. We fitted a Generalized Linear Model (GLM) with the number of trials per subject and week in which mild signs of stress were recorded as the response variable, and season of testing included as an exploratory categorical predictor, assuming a Poisson distribution for the response.

### Physical problem-solving

For the performance in the physical problem-solving task we used the number of problems solved over the number of problems available at each developmental timepoint as the response variable, assuming a binomial distribution, and as explanatory variables we used (a) the average time of exploration of the test apparatus for all trials during each week, (b) sex, and (c) week.

### Role of memory

To compare the performance of subjects first presented with the problem-solving task at ten weeks of age with that of subjects first presented at seven weeks and re-presented at ten weeks, we fitted a GLMM with the proportion of problems solved out of those available as the response, assuming binomial distribution. Explanatory variables included sex, the average time spent investigating the apparatus across all trials at each developmental time point, and week as a factor with three levels according to the experimental condition: (1) first presentation at seven weeks of age, (2) first presentation at ten weeks of age, and (3) second presentation at ten weeks of age. We included individual identity as a random factor.

### Development of individual consistency in task performance

To investigate whether individual consistency changed over the course of early life, we used a multivariate hierarchical modelling approach, using the brms package (61). For each cognitive test, we fitted 5 Generalized Linear Models, one for each week of testing. We fitted the same set of fixed response, explanatory variables and assumed distribution of the models used to investigate individual performance over development, with the exclusion of week of age. For the random part, we allowed the intercept calculation to vary for individual by fitting individual random intercepts. Then, for each cognitive test, we used the multivariate hierarchical design to correlate individual intercepts calculated at each week with each other. Random individual intercepts were calculated as deviation from the global intercept, which for each model was the specific week investigated. Thus, we obtained within each week a set of individual values that highlighted whether the subject performed above or below average, while factoring out the covariates of interests that could confound cognitive performance and create spurious correlations. Thus, comparing individual intercepts among weeks allowed us to investigate whether subjects that performed above average at one point in time (e.g. week 10) were more likely to perform above average at a future developmental stage (e.g. week 13).

### Correlation among task performance of the three cognitive traits

#### Over the developmental period

To investigate whether performance in one cognitive task was related to performance in another, we extended our hierarchical models into a multivariate framework. We refitted the models described in the section “Population-level developmental trajectories and individual variation”, retaining for the spatial cognition task the version including study design as a predictor, as it outperformed the alternative model with the binary ‘correct exit changed’ variable. This multivariate approach allowed us to address two questions: (1) whether subjects who performed better than average in one task also performed better in another, assessed by comparing individual intercepts across tasks/models; and (2) whether subjects who improved more rapidly than average in one task showed similar improvement in another, assessed by comparing individual slopes for the effect of week on performance across tasks/models.

#### At specific developmental time points

To investigate whether relationships between performance in any pair of cognitive tasks changed over the course of development, we again used a multivariate hierarchical modelling approach. For each week of testing, we fitted three models (one for each cognitive task) using the same specification (response, predictors, random effects, and assumed distributions) as described in the section “Development of individual consistency in task performance”. This framework allowed us to assess, for each week, whether subjects performed above or below average after accounting for confounding variables (individual intercepts). By comparing performance across tasks, we then determined whether relationships between cognitive traits varied across early life.

## Results

We report model estimates as the mean with the 95% credible interval in square brackets. As a measure of evidence, we calculated the proportion of posterior samples supporting the estimated effect (PP_support_). We encourage readers to treat PP_support_ as a continuous measure of evidence without clear boundaries, and to make their own decision on sufficient levels of credibility. However, to ease interpretation and discussion, we considered effects with PP_support_ ≥ 95% as supported, and effects with PP_support_ of between 70% and 95% as suggestive but inconclusive trends (62). We report the proportion of remaining variation (i.e., not explained by fixed effects) explained by random effects as the median with the 95% credible interval in square brackets. Although negative values for this proportion are numerically impossible, they can appear as artefacts of high variability in the posterior distribution (60). When the lower bound of the 95% credible interval extended below zero, we truncated it at zero and interpreted the effect as not being convincingly different from zero.

### Population-level developmental trajectories and individual variation

#### Inhibition control

We found support for an increase in performance in the inhibitory control task over successive developmental week (mean effect of week = 0.09, 95% CI [0.05, 0.13], PP_support_ = 100%, Figure 2). There was no evidence that males and females differed in their probability of passing the inhibitory control test (effect of males, relative to females = 0.08, [-0.55, 0.74], PP_support_ = 60.6%).

Individual differences in learning curves (random slopes) accounted for 11% [0%, 45%] of the remaining variability that was not explained by fixed effects (henceforth: ‘remaining variability’), while individual differences in baseline performance (random intercepts) accounted for 28% [0%, 56%], and the combination of the two explained 18% [0%, 44%]. Thus, we find little evidence for individual deviation from population-level developmental trends in this cognitive task. Individual baseline performance without the effect of covariates (random intercepts) and learning curves (random slopes) showed a non-supported trend towards a negative correlation (correlation = -0.46, [-0.95, 0.62], PP_support_= 83.0%).

#### Spatial cognition

We found support for increasing performance in the spatial cognition task with successive developmental week, with subjects making less errors later in their development (week = - 0.04, [-0.07, -0.01], PP_support_ = 99.8%, Figure 2). Performance was also influenced by stress, with subjects making more errors when showing mild signs of stress (effect of stress = 0.36, [0.22, 0.50], PP_support_ = 100%). We found no support for a difference between males and females in the average number of errors per trial (effect of male, relative to females = -0.06, [-0.64, 0.53], PP_support_ = 58.1%). When the correct exit was kept the same from the previous testing week, we found a non-supported trend in favour of subjects making less errors when the exit was consistent among consecutive weeks (effect of consistent exit, relative to inconsistent = -0.09, [-0.38, 0.19], PP_support_= 72.6%). Test design influenced performance, as we found that subjects made less errors during the second season (effect of second season, relative to first season = -1.01, [-1.73, -0.22], PP_support_= 99.2%), while at the same time more likely to show mild signs of stress (second season, relative to first season = 1.25, [0.02, 2.76], PP_support_= 97.6%). Accordingly, we used the model including the study design to calculate the proportion of remaining variance explained by random factors, to factor out possible spurious individual effects due to the two seasons.

Including individual learning curves (random slopes) in our model for the development of spatial cognition accounted for 72% [32%, 91%] of remaining variation, while including individual baseline performances (random intercepts) accounted for 80% [24%, 95%] of it. Including the combination of individual learning curves and baseline performance in the model accounted for 51% [0%, 81%] of remaining variability. Thus, we find strong support for individual differences in the development of performance in the spatial cognition task in both baseline abilities and learning curves. The proportion of remaining variability was lower when random slopes and random intercepts were included together, possibly due to the covariance between the two factors (correlation = -0.69, [-0.92, -0.26], PP_support_= 99.6%). This suggests that when analysed separately, each factor also captures part of the variability explained by the other. The negative covariance between random slopes and intercepts indicates that individuals with above-average baseline performance (i.e., making more errors after accounting for covariates) tended to show below-average rates of developmental change (i.e., a faster decrease in errors over time). Thus, subjects that performed worse at a baseline showed faster learning curves during development.

#### Physical problem-solving

The performance in the physical problem-solving task increased over ontogeny, with the effect of developmental week having a positive effect (week = 0.26, [0.17 - 0.36], PP_support_= 100%, Figure 2). Subjects that spent more time exploring and interacting with the apparatus were more likely to solve the problems it posed (exploration time = 0.03, [0.01, 0.05], PP_support_ = 100%). We found a suggestive trend for males being less likely to remove an obstacle than females, though we did not find full support for this finding (males, relative to females = -0.42, [-1.35, 0.51], PP_support_ = 82.3%). Individual learning curves (random slopes) accounted for 60% [43%, 72%] of the remaining variability, while individual baseline performances (random intercepts) accounted for 44% [12%, 66%] of it. The combination of random slopes and intercepts accounted for 44% [21%, 63%] of remaining variability. Thus, we find moderate to medium to high levels of support for differences in both individual baseline performances and individual rates of development (learning curves). The decrease in variability explained when both random slopes and intercepts were included is possibly due to the two random factors covarying with each other (correlation = -0.65, [-0.88, -0.25], PP_support_= 99.6%) and explaining in isolation part of the other factor’s explained variability. In addition, this correlation highlights that subjects with poorer baseline performance showed faster learning curves during development.

#### Role of memory

Subjects who were presented with the physical problem-solving task for the first time at seven weeks of age performed worse than subjects who were presented with it for the first time at ten weeks of age (first presentation at seven weeks, relative to first presentation at ten weeks = -1.07, [-2.17, 0.02], PP_support_ = 97.2%). We did not find support for a difference in performance between subjects who interacted with the apparatus for the first time at ten weeks and those that interacted with it for the second time at ten weeks, although there was a non-supported trend for the latter outperforming the former (second presentation at ten week, relative to first presentation at ten week = 0.35, [-0.75, 1.48], PP_support_ = 73.4%).

### Development of individual consistency in task performance

Detailed model outputs can be found in Supplementary Table S4.

#### Inhibitory control

We did not find support for individual performance at one developmental time point to predict performance at the subsequent time point (Figure 3). A non-supported positive trend was observed between week 7 and week 10 which disappeared in later comparisons. Likewise, performance at the onset of sub-adulthood (week 24) was not predicted by any earlier time point (Figure 3).

**Figure 3:**
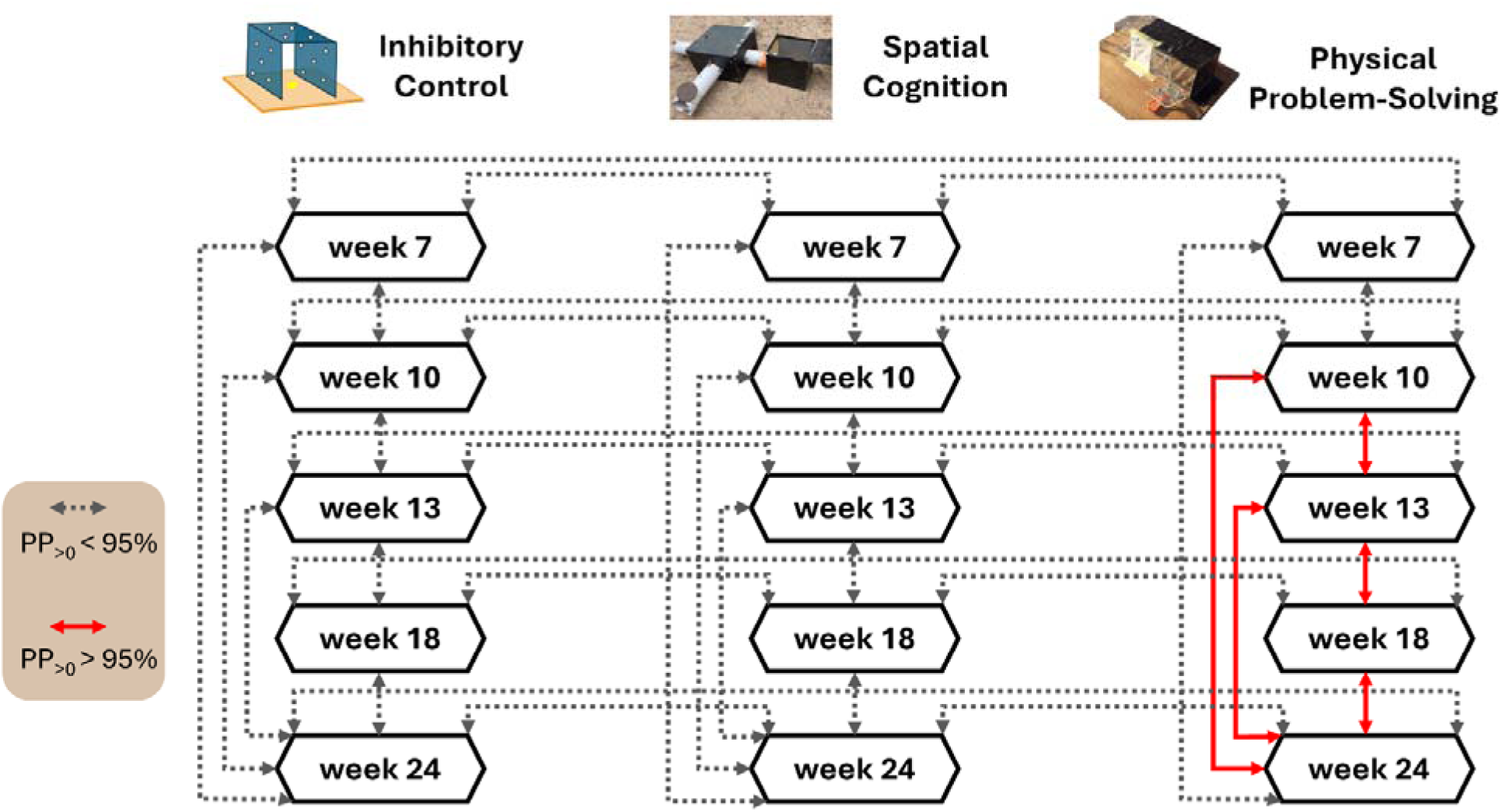
Summary of correlations between individual random intercepts among tasks, and between tasks, over the developmental period. Full, red lines are positive, supported correlations, while dotted lines are non-supported correlations.

#### Spatial cognition

We did not find support for consistency in individual performance across developmental time points (Figure 3), although correlation estimates were consistently positive. A non-supported positive trend was observed between week 7 and week 10 and between week 13 and week 18, while support for the same correlation at week 10–13 was even weaker. Likewise, performance at the onset of sub-adulthood (week 24) could not be convincingly predicted by earlier time points.

#### Physical problem-solving

We found strong support for consistency in individual performance across consecutive developmental weeks, emerging early in ontogeny (Figure 3). A non-support positive trend was present between week 7 and week 10, which became supported in week comparisons from week 10 onwards. Reflecting this pattern, performance at the onset of sub-adulthood (week 24) was predicted by performance at week 18, 13 and 10, while at the earliest stage (week 7) the relationship remained a non-supported positive trend.

### Correlation among task performance of the three cognitive traits over the developmental period

Because performance metrics were scaled in opposite directions (higher values indicated better performance in inhibitory control and physical problem-solving, while lower values indicated better performance in spatial cognition) we mirrored the correlation estimates involving spatial cognition. Accordingly, positive correlations between task performances consistently indicate that better performance in one task was associated with better performance in the other.

#### Inhibitory control & spatial cognition

We found a non-supported trend between the rates of development of the two traits, calculated as the random slope of the effect of week on performance for each subject, with faster learning curves in inhibitory control performance associated with slower learning curves in spatial cognition performance (correlation = -0.21, [-0.83, 0.43], PP_support_ = 73.7%). When comparing individual baseline performances after removing the effects of covariates (random intercepts), we found another non-supported trend, with a better performance in inhibitory control associated with a worse performance in spatial cognition (correlation = -0.36, [–0.81, 0.13], PP_support_ = 91%).

#### Spatial cognition & physical problem solving

No supported correlation was detected between developmental rates (correlation = -0.02, [–0.47, 0.41], PP_support_ = 52.6%). At the baseline performance level, we observed a non-supported trend, with a better performance in the physical cognition task associated with a worse performance in the spatial cognition task (correlation = -0.27, [–0.70, 0.16], PP_support_ = 88.4%).

#### Inhibitory control & physical problem solving

We found non-supported positive trends for both developmental rates (correlation = 0.21, [–0.43, 0.80], PP_support_ = 75.7%) and baseline individual performance (correlation = 0.22, [–0.30, 0.72], PP_support_ = 78.8%), with better inhibitory control performance associated with better physical problem-solving performance.

### Correlation among task performance of the three cognitive traits at specific developmental time points

Similarly to the previous section, we mirrored the correlation estimates involving spatial cognition so that positive correlations between task performances consistently indicate better performance in one task being associated with better performance in another. Detailed model outputs can be found in Supplementary Table S5.

#### Inhibitory control & spatial cognition

We found no supported associations between inhibitory control and spatial cognition at any time point (Figure 3). A non-supported negative trend was present at 7 weeks of age.

#### Spatial cognition & physical problem solving

Associations between spatial cognition and physical problem solving also lacked support at all time points (Figure 3). Estimates were consistently positive across development, with a non-supported positive trend at weeks 10 and 13.

#### Inhibitory control & physical problem solving

No supported associations or unsupported trends were detected between inhibitory control and physical problem solving at any time point (Figure 3).

## Discussion

In this study, we investigated the developmental trajectories of three ecologically relevant cognitive traits in wild meerkats. We found evidence that cognitive performance increases in meerkats over ontogeny, spanning from early life to early sub-adulthood. Cognitive development was present for all measured traits: inhibition control, spatial cognition and physical problem-solving skills. At the population level, inhibitory control and spatial cognition (spatial orientation and memory) showed a linear increase in performance and did not reach a plateau within the developmental timeframe investigated, while physical problem-solving skills showed the fastest increase in performance (steepest learning curves) and reached a plateau before subjects grew into sub-adulthood (Figure 2). We found substantial evidence for individual variation in the development of both spatial cognition and physical problem-solving ability but not in the development of inhibitory control, suggesting the latter to be under developmental constrain. Of all cognitive traits, we found support for individual consistency in performance only in physical problem-solving ability, which arose early and endured until the end of the developmental period studied. Contrary to our expectations based on male dispersal and the cognitive requirements of interactions with outgroup conspecifics, we did not find evidence for differences in cognitive performance between males and females in any of the three traits investigated. Furthermore, although we expected cognitive performance to positively covary in meerkats due to their extensive reliance on social learning during development (42), we found no support for such covariance during early development, either in individual trajectories or at discrete time points.

The timing of cognitive development in each trait can be linked to specific socio-ecological demands in meerkats. Early in their ontogeny meerkats must reach sufficient independent foraging competence, as social provisioning by adults stops around the twelfth week of age. This timing aligns with our findings on the cognitive ability to solve extractive physical problems, where meerkats had a 76% (CI 95% [57%, 89%]) probability of solving a physical barrier assuming 30 seconds of exploration at 12 weeks of age. Despite an early onset of independent foraging ability (50), meerkats remain within their family groups way beyond when their foraging competence is reached (25) and can rely on social information and cues from their group members for spatial navigation and coordination for a longer period. Before nutritional independence, pups can react to alarm calls by following the heading of nearby older individuals to reach safety (30,41). Afterwards, decisions about group movement are mostly influenced by older individuals (63). Accordingly, our findings suggest that the development of spatial cognition is slower and lacks a clear plateau within the developmental timeframe we investigated (Figure 2), suggesting that this cognitive trait is less critical early in life. Since investment in neurological tissue is expensive (9,11,44,64), partly relying on collective information may be an energetically efficient strategy (65), especially early in life when demands on independent spatial memory are lower. Similarly, there are lower demands for inhibitory control early in meerkat life, as juveniles are seen to contribute less to cooperative activities than older individuals (25). Selection may even favour low inhibitory control early in pup development, as adults react to signals of hunger and feed pups that beg more intensively and persistently more often (66,67). However, these benefits progressively decline as pups grow, because adults stop responding to their begging calls when they reach three to four months of age (68), and adults may need to reach a minimum baseline of inhibition ability for effective cooperative activities. Accordingly, we found that inhibitory control shows a faster development than spatial cognition but developed more slowly than physical problem-solving and lacks a clear plateau before sub-adulthood. Thus, life-history demands likely influence the developmental timings of inhibitory control, spatial cognition, and physical problem solving in meerkats.

We found little individual variation in inhibition control for both baseline performance (without the effect of sex and development) and developmental rates (learning curves). In addition, we did not find evidence for a difference in performance between males and females. We hypothesized that the ability to inhibit behaviour would be a cognitive underpinning of cooperative behaviours (26,45). Although meerkats show consistent and repeatable differences in their contributions to cooperative behaviours (69,70) and males’ relative contributions differ from females’ (25), we did not find evidence that differences in the early development of inhibitory control could underlie these patterns. Instead, the ontogeny of this cognitive trait seems developmentally canalized, with limited influence of environmental or genetic factors (71). Inhibitory control is still likely to play a part in cooperative activities, because adult meerkats need a minimum level of inhibitory control to perform cooperative activities by supressing self-serving behaviours such as ingesting foraged prey, after which internal and physiological states are likely to drive individual variation. Specifically, in meerkats, contributions to cooperative activities are linked to body condition (72), foraging success (73), and individual necessities (e.g., gathering information for future dispersal) (25), and are under hormonal control (74). The conserved early life development of inhibitory control in meerkats could also result from low genetic variability underlying this trait, arising from selection removing sub-optimal variance. However, although individual variation was not evident during early development, it may arise later in response to ecological or social pressures. Future studies should investigate if adult performance is repeatable and whether it is linked to enduring differences in cooperative contributions.

In the spatial cognition task, individuals differed in both baseline performances (without the effect of development, sex, apparatus design or stress), and rates of development (learning curves). In addition, although males are the dispersing sex, they performed similarly to females. Individual differences in baseline performances could be due to intrinsic cognitive characteristics or spatial orientation strategy. Spatial orientation can be either based on position in space in relation to an individual’s own position and direction (egocentric orientation), or on external cues like landmarks (allocentric orientation) (75). Our study design focused on egocentric orientation, as the apparatus was opaque and the orientation of the correct exit in relation to any other perceivable external cue like magnetic field and group position (through sound) was inconsistent among trials. We found that when the apparatus was even partially transparent during the first season, meerkats were more likely to make errors, which was not explained by stress levels nor other study design decisions. Therefore, a decrease in performance during the first season is likely linked to the partial transparency of the apparatus, and consequently individual spatial orientation strategy. In addition, the ability to inhibit intuitive responses could have also influenced performance in a partially transparent apparatus, as subjects had to inhibit their attempts to go through a transparent obstacle. However, we did not find that performance in the inhibition control predict performance in the spatial maze in our dataset. Interestingly, individuals that were below average in baseline performances (without the effect of development) had faster learning curves, suggesting that there are minimum levels of cognitive functions that individuals are selected to reach at the onset of sub-adulthood.

In the physical problem-solving task, we found individual differences in both baseline performances (without the effect of development, sex and explorative effort) and rates of development. Individual performances were also consistent from the tenth week until sub-adulthood, suggesting that differences in physical problem-solving performance in meerkats arise early in development and are enduring. Individuals who were presented the task for the first time at ten weeks of age outperformed their peers who were presented it for the first time at seven weeks and were not substantially different from those same peers when the latter were re-presented with it at ten weeks of age. This suggest that the increase in performance we observe during early development is not exclusively due to memory and previous test experience, but in fact likely linked to the development of brain functions and motor skills. Although motor skills are usually seen as a confounding factors when investigating underlying cognitive ability, recent work has questioned whether motor skills and cognitive abilities could be meaningfully disentangled from each other (76). As such, a physical problem-solving test where motor skills are involved is a meaningful assessment of cognitive ability. Our results can be linked to different meerkat foraging strategies. Individuals either scan the ground and find preys on the surface or identify specific places in the environment where they dig up to thirty centimetres for the chance of finding prey below ground. Foraging strategy is known to change with age (33,77), and young individuals may also vary in their tendency to engage in energetically costly foraging with uncertain returns but valuable learning opportunities. These different strategies could reflect variation in learning curves in their physical problem-solving abilities and be linked to socio-ecological conditions experienced during development. Similarly to performance in the spatial cognition task, when individuals had a below average baseline performances, they showed faster learning curves. Thus, selection may shape meerkats to reach a minimal necessary level of physical cognition at the onset of sub-adulthood to handle ecological challenges arising from independent foraging.

Individual performances across the three tasks were uncorrelated, both in their learning curves, individuals’ baseline abilities, or performance at any developmental time point right up to sub-adulthood, suggesting that in meerkats inhibitory control, spatial cognition (navigation and memory) and problem-solving ability are likely modular rather than domain-general adaptations (42). Their cognitive underpinnings may therefore be independent of one another, and potentially subject to separate selective pressures, at least during early development. Moreover, we found that meerkats’ chances of successfully passing a standardized inhibitory control test at the onset of sub-adulthood (model predicted performance = 53.4%, 95% CI [37.8%, 68.1%]) were worse than 20 of 32 species investigated by MacLean and colleagues (51). However, meerkat performance in inhibitory control may not be fully developed by this point in their ontogeny. Altogether, our results from a cooperative mammal do not align with hypotheses that link social learning (42) or cooperative care (26,44,45) to the evolution of general intelligence or enhanced cognitive abilities in primates. As such, our study highlight how selective pressures may not be operating uniformly and require fine scale developmental investigations including different animal taxa.

## Conclusion

We present the first detailed longitudinal study on cognitive development in a carnivorous mammal under natural conditions. We provide evidence that three cognitive traits (inhibition control, spatial cognition, and physical problem-solving) emerge at different times of early development in meerkats, likely because of distinct socio-ecological demands. By investigating individual deviations from population-level trends, we found that inhibitory control follows a conserved developmental trajectory, while problem-solving ability and spatial cognition show inter-individual developmental variability and are likely to be influenced by socio-ecological or genetic factors. Cognitive performance across the three cognitive traits did not correlate within individuals, either at singular points in time or in their developmental trajectories, suggesting modular rather than domain-general cognitive underpinnings. These findings highlight the importance of studying cognitive evolution across different taxa, social systems and developmental timescales. Future studies should uncover the sources of the individual variation in developmental trajectories and their adaptive consequences.

### Ethical statement

This research was approved by the University of Pretoria (Ethics Permit NAS061/2022) and the Northern Cape Province Department of Environment and Nature Conservation, South Africa, (FAUNA 0931/2022).

## Acknowledgements

We thank the Swiss National Science Foundation for funding this work through the grant no. PZ00P3_202052 awarded to SF. We thank the Kalahari Research Trust and the Kalahari Meerkat Project for organisational support and access to the study population. We are grateful to neighbouring landowners for permission to use their land. We are particularly thankful to Marta Manser for discussion and feedback on the manuscript. We also thank Walter Jubber, Jordan Martin, Josh Arbon, and Emily Stott for their valuable input and support. Finally, we appreciate all members of the Kalahari Research Centre (staff, volunteers, managers, and researchers) for maintaining the research infrastructure, habituating and monitoring the meerkats, and providing professional and logistical support during fieldwork.

## Declaration of generative AI and AI-assisted technologies in the writing process

During the preparation of this work the authors used ChatGPT (OpenAI, 2025) to improve grammar, clarity and quality of language. After using this tool, the authors reviewed and edited the content as needed and take full responsibility for the content of the published article.

## Author contributions

T.S.: Conceptualization, Data Curation, Formal Analysis, Investigation, Methodology, Software, Visualization, Writing – Original Draft. E.P.: Investigation, Data Curation, Writing – Review & Editing. A.V.J.: Methodology, Supervision, Writing – Review & Editing. S.I.F.F.: Conceptualization, Methodology, Supervision, Funding Acquisition, Writing – Review & Editing.

