## Supplementary Table S4 for "Developmental trajectories of cognitive traits in meerkats match socio-ecological demands"

**Table S4:** Model outputs investigating the development of individual consistency in task performance across ontogeny.

| Cognitive Trait |  | Variable 1 | Variable 2 | Correlation | Lower 95% CI | Higher 95% CI | PP_support_ |
| --- | --- | --- | --- | --- | --- | --- | --- |
| Inhibitory Control |  | Week 7 | Week 24 | 0.00 | -0.67 | 0.65 | 49.6% |
|  |  | Week 10 | Week 24 | 0.05 | -0.63 | 0.67 | 56.7% |
|  |  | Week 13 | Week 24 | -0.02 | -0.67 | 0.64 | 52.5% |
|  |  | Week 18 | Week 24 | 0.02 | -0.65 | 0.67 | 53.3% |
|  |  | Week 7 | Week 10 | 0.22 | -0.43 | 0.80 | 75.2%^∆^ |
|  |  | Week 10 | Week 13 | -0.07 | -0.72 | 0.55 | 58.0% |
|  |  | Week 13 | Week 18 | -0.09 | -0.74 | 0.58 | 60.4% |
| Spatial Cognition |  | Week 7 | Week 24 | 0.19 | -0.56 | 0.88 | 69.4% |
|  |  | Week 10 | Week 24 | 0.16 | -0.60 | 0.83 | 65.9% |
|  |  | Week 13 | Week 24 | 0.13 | -0.61 | 0.86 | 62.8% |
|  |  | Week 18 | Week 24 | 0.18 | -0.58 | 0.87 | 68.4% |
|  |  | Week 7 | Week 10 | 0.29 | -0.33 | 0.85 | 80.2% |
|  |  | Week 10 | Week 13 | 0.17 | -0.52 | 0.82 | 67.6% |
|  |  | Week 13 | Week 18 | 0.27 | -0.42 | 0.89 | 77.1%^∆^ |
| Physical Problem-Solving |  | Week 7 | Week 24 | 0.24 | -0.29 | 0.76 | 80.6%^∆^ |
|  |  | Week 10 | Week 24 | 0.51 | 0.05 | 0.92 | 97.1%* |
|  |  | Week 13 | Week 24 | 0.48 | 0.05 | 0.88 | 97.2%* |
|  |  | Week 18 | Week 24 | 0.59 | 0.14 | 0.96 | 98.3%* |
|  |  | Week 7 | Week 10 | 0.27 | -0.28 | 0.80 | 81.8%^∆^ |
|  |  | Week 10 | Week 13 | 0.69 | 0.33 | 0.98 | 99.7%* |
|  |  | Week 13 | Week 18 | 0.68 | 0.32 | 0.98 | 99.8%* |

*Note:* For each cognitive trait, Variable 1 and Variable 2 represent performance at the respective developmental weeks. Individual cognitive performances were calculated as random intercepts after controlling for confounding variables (see main text). For correlations involving spatial cognition, the sign was reversed to ensure that positive correlations consistently represent better performance across traits. Correlations reflect the posterior mean with associated 95% credible intervals. PP_support_ indicates the proportion of posterior samples supporting the estimated correlation. Asterisks(*) indicate supported effects (PP_support_ ≥ 95%). Triangles(^∆^) indicate unsupported trends (PP_support_ 75%-95%).
