## Supplementary Table S5 for "Developmental trajectories of cognitive traits in meerkats match socio-ecological demands"

**Table S5:** Model outputs investigating the development of correlation among individual performance in three cognitive traits across ontogeny.

| Week | Variable 1 | Variable 2 | Corr. | Lower 95% CI | Higher 95% CI | PP_support_ |
| --- | --- | --- | --- | --- | --- | --- |
| 7 | Spatial Cognition | Inhibitory Control | -0.17 | -0.77 | 0.43 | 71.4% |
|  | Spatial Cognition | Physical Problem-Solving | 0.03 | -0.46 | 0.51 | 54.2% |
|  | Inhibitory Control | Physical Problem-Solving | 0.01 | -0.64 | 0.64 | 51.6% |
| 10 | Spatial Cognition | Inhibitory Control | -0.39 | -0.81 | 0.07 | 94.3%^∆^ |
|  | Spatial Cognition | Physical Problem-Solving | 0.23 | -0.18 | 0.63 | 85.2%^∆^ |
|  | Inhibitory Control | Physical Problem-Solving | 0.01 | -0.47 | 0.46 | 51.0% |
| 13 | Spatial Cognition | Inhibitory Control | -0.02 | -0.73 | 0.71 | 52.9% |
|  | Spatial Cognition | Physical Problem-Solving | 0.44 | -0.09 | 0.89 | 93.6%^∆^ |
|  | Inhibitory Control | Physical Problem-Solving | -0.06 | -0.71 | 0.61 | 57.1% |
| 18 | Spatial Cognition | Inhibitory Control | -0.12 | -0.81 | 0.56 | 63.6% |
|  | Spatial Cognition | Physical Problem-Solving | 0.03 | -0.50 | 0.54 | 53.9% |
|  | Inhibitory Control | Physical Problem-Solving | -0.08 | -0.75 | 0.62 | 59.1% |
| 24 | Spatial Cognition | Inhibitory Control | 0.06 | -0.69 | 0.80 | 55.6% |
|  | Spatial Cognition | Physical Problem-Solving | 0.14 | -0.59 | 0.82 | 65.4% |
|  | Inhibitory Control | Physical Problem-Solving | 0.21 | -0.51 | 0.91 | 71.6% |

*Note:* For each cognitive trait, Variable 1 and Variable 2 represent performance in the respective cognitive trait. Individual cognitive performances were calculated as random intercepts after controlling for confounding variables (see main text). Correlations (“Corr.”) reflect the posterior mean with associated 95% credible intervals. For correlations involving spatial cognition, the sign was reversed to ensure that positive correlations consistently represent better performance across traits. PP_support_ indicates the proportion of posterior samples supporting the estimated correlation. Asterisks(*) indicate supported effects (PP_support_ ≥ 95%). Triangles(^∆^) indicate unsupported trends (PP_support_ 75%-95%).
