## Supplementary Text S2 for "Developmental trajectories of cognitive traits in meerkats match socio-ecological demands"

**Text S2:** Extended details on habituation procedure for inhibitory control, spatial cognition, and physical problem-solving, as well as apparatus design for the physical problem-solving task.

**Inhibitory control**

*Habituation procedure:* At the start of each testing week, we habituated subjects by presenting an opaque version of the test apparatus. The apparatus was presented to the subjects in the same way as the test apparatus, with the barrier perpendicular to the subject. To confirm that subjects had learned the rule associated with the apparatus, each subject was presented with the opaque version until it reached a criterion of four successful trials out of five. A trial was considered successful when the meerkat detoured to the open side and retrieved the food reward without touching or sniffing the opaque panel and without pausing in front of it for longer than two seconds.

**Spatial cognition**

*Habituation procedure:* Individuals were habituated to the apparatus at each developmental week before testing by positioning them in the apparatus while all three exit arms were fully open for multiple times to ensure they were comfortable with being inside the box and the arms. During habituation, we let each subject (at each developmental week) exit the fully open apparatus from at least two different arms, to avoid them making associations to a specific exit before actual testing.

**Physical problem-solving**

*Test design:* The apparatus included four contiguous chambers, each containing a food reward inside. The sides and top of the apparatus were opaque, but the division between internal chambers was made of transparent plastic. Three chambers were covered by a different obstacle to be removed or opened, with one additional chamber left unobstructed to maintain motivation to participate. One problem was a sheet of toilet paper that was taped to fully close the chamber. The toilet paper for the 7^th^ week of testing was single plied and for successive weeks was double plied, to adjust for the lowest physical strength of seven weeks young pups. The second problem was a transparent flap hanging from the top of the chamber, that needed to be pushed to access the food inside. The third problem featured a transparent door with a hinge on the lower side of the chamber that needed to be pulled opened.

*Habituation procedure:* Each week before testing subjects were presented with an apparatus with the four food-baited chambers without any obstacle to be removed. We only started testing once a subject had eaten from all chambers once in at least two separate presentations to ensure familiarization with the apparatus and motivation to participate in the cognitive trials.
