## Supplementary Table S1 for "Developmental trajectories of cognitive traits in meerkats match socio-ecological demands"

**Table S1:** Summary of deviations from the planned data collection. Cognitive traits are inhibitory control (IC), spatial cognition (Maze), and physical problem-solving (Problem). Deviations are categorised by the number of trials affected. Planned data collection consisted of five IC trials, four Problem trials, and four Maze trials per individual per developmental week (7, 10, 13, 18 and 24).

| **ID** | **Week** | **Cognitive trait** | **Type of deviation** | **Issue** |
| --- | --- | --- | --- | --- |
| VALF008 | 7 | IC | single trial | 5th trial missing recording |
| VDDM006 | 7 | IC | single trial | 4th trial missing recording |
| VJXF190 | 7 | IC | single trial | 5th trial excluded (not a straight approach) |
| VALM016 | 7 | Maze | single trial | 4th trial excluded because correct exit was stuck |
| VJXF189 | 7 | Maze | single trial | 4th trial missing due to miscount |
| VUBM073 | 7 | Maze | single trial | 2nd trial was considered a mistrial |
| VUBF071 | 10 | IC | single trial | 3rd trial excluded (individual failed but did not access food reward) |
| VALF008 | 10 | Problem | single trial | Presented 5 times (mistaken for VALM009) |
| VALM009 | 10 | Problem | single trial | Presented 3 times (mistaken for VALF008) |
| VJXF190 | 10 | Problem | single trial | 2nd trial corrupted recording |
| VDDM007 | 10 | Problem | single trial | Presented 5 times (mistaken ID) |
| VUBF071 | 10 | Maze | single trial | 4th trial missing (field miscount) |
| VSIM020 | 10 | Maze | single trial | 2nd trial was considered a mistrial |
| VDDM007 | 13 | IC | single trial | 1st trial missing recording |
| VLF297 | 13 | IC | single trial | Presented only 4 times |
| VLF299 | 13 | IC | single trial | Presented only 4 times |
| VLM298 | 13 | IC | single trial | Presented only 4 times |
| VLM300 | 13 | IC | single trial | Presented only 4 times |
| VALM019 | 18 | IC | single trial | 1st trial excluded for social disturbance |
| VLF299 | 18 | Problem | single trial | 3rd trial missing recording |
| VLF297 | 24 | IC | single trial | Presented 4 times because mistaken for VLM298 |
| VLM298 | 24 | IC | single trial | Presented 6 times because mistaken for VLF297 |
| VECM055 | 7 | Maze | multiple trials | Single video for all trials corrupted |
| VUBM073 | 13 | IC | multiple trials | 4^th^, 5th trials excluded (individual failed but did not access food reward) |
| VALM016 | 13 | Maze | multiple trials | Missing recording all trials |
| VUBF071 | 18 | IC, Problem | multiple trials | Data could not be collected |
| VUBM072 | 18 | IC, Problem | multiple trials | Data could not be collected |
| VUBM073 | 18 | IC, Problem | multiple trials | Data could not be collected |
| VALF017 | 18 | Maze | multiple trials | Single video for all trials corrupted |
| VLF297 | 18 | Maze | multiple trials | Data could not be collected |
| VLM300 | 18 | Maze | multiple trials | Data could not be collected |
| VSIM020 | 18 | Maze | multiple trials | Data could not be collected |
| VSIM021 | 18 | Maze | multiple trials | Data could not be collected |
| VSIF023 | 18 | Maze | multiple trials | Data could not be collected |
| VSIM024 | 18 | Maze | multiple trials | Data could not be collected |
| VSIM020 | 24 | IC, Maze, Problem | multiple trials | Data could not be collected |
| VSIM021 | 24 | IC, Maze, Problem | multiple trials | Data could not be collected |
| VSIF023 | 24 | IC, Maze, Problem | multiple trials | Data could not be collected |
| VSIM024 | 24 | IC, Maze, Problem | multiple trials | Data could not be collected |
| VUBF071 | 24 | IC, Maze, Problem | multiple trials | Data could not be collected |
| VUBM072 | 24 | IC, Maze, Problem | multiple trials | Data could not be collected |
| VUBM073 | 24 | IC, Maze, Problem | multiple trials | Data could not be collected |
| VJXF188 | 24 | IC, Maze, Problem | multiple trials | Data could not be collected |
| VJXF189 | 24 | IC, Maze, Problem | multiple trials | Data could not be collected |
| VJXF190 | 24 | IC, Maze, Problem | multiple trials | Data could not be collected |
| VJXF191 | 24 | IC, Maze, Problem | multiple trials | Data could not be collected |
