## Supplementary Table S3 for "Developmental trajectories of cognitive traits in meerkats match socio-ecological demands"

**Table S3.** List of definitions for the scoring of cognitive trials.

| **Cognitive Trait** | **Scored Behaviour** | **Explanation** |
| --- | --- | --- |
| Physical problem-solving | Explore | Any attempt to sniff, smell, bite, dig at the apparatus or look at the food reward (egg) inside the chambers and follow it with its gaze while moving, or physically/visually exploring the sides, roof or doors of the apparatus. |
| Physical problem-solving | Solve Obstacle 1 (Toilet Paper) | The task is solved when the animal access the food inside or a hole in the toilet paper is open/big enough for the animal to enter, permanent, and the meerkat interacts with it; the latter usually result in the meerkat eating the food reward, but the problem is scored as solved even if the meerkat does not do consume the reward due to external influences (e.g., refuses the reward or runs away because of a conspecific alarm). |
| Physical problem-solving | Solve Obstacle 2 (Catflap) | The task is solved when the meerkat’s head is inside enough to access the reward or if the individual grabs the reward from inside by pulling it out with its paws. |
| Physical problem-solving | Solve Obstacle 3 (Pull Door) | The task is solved when the door is open for the animal to retrieve the reward. (Accident) If the individual has access or solves the pull door while bumping accidentally into the apparatus or using a motor action that was not aimed at the pull door chamber. If the individual uses any motor action towards the pull door and the pull door opens “too easily” because of the angle of the test, that is not considered a mistake. |
| Physical problem-solving | Test over | A trial is over when (a) All three problems have been solved, and the apparatus leaves the ground, (b) approximately 60 seconds or more have passed and is ended by the experimenter, (c) the meerkat loses interest in the apparatus, (d) due to external influences, (e) the meerkat runs away because of a conspecific alarm. |
| Spatial cognition | Start trial | When the individual is first inside the starting box. |
| Spatial cognition | Position | At any point from the start of the trial (“start trial”) until the end of the trial (“end trial”), note the position of the individual within the maze. An individual is considered in a location if more than half of its body (excluding the tail) is within one location. The locations are “Starting Box / Arm”, “Main Arena” and each arm (Centre, Right and Left, with reference to the starting box / arm). |
| Spatial cognition | Stress | (Yes) if the meerkat made distress calls for at least 3 seconds consecutively at any point during the testing. |
| Spatial cognition | End trial | The trial ends when the end is called or the meerkat was observed exit the correct arm This is scored even if the meerkat finds the exit (e.g., clearly peaks out of the correct end arm) but does not fully exit and returns to the main arena. |
| Inhibitory control | Result | A success was defined when the subject obtained the food reward without making physical contact with the transparent sides of the apparatus. A failure was defined as any contact with the transparent sides, regardless with which body part. |
